# Preliminary report on the molecular epidemiology of mpox, South Africa 2024

**DOI:** 10.1101/2024.11.07.622575

**Authors:** Jonathan Featherston, Antoinette Ann Grobbelaar, Allison Glass, Dominique Goedhals, Kim Hoek, Jean Maritz, Kamini Govender, Nevashan Govender, Arshad Ismail, Chantel le Roux, Terry Marshall, Daniel Morobadi, Mishalan Moodley, Naazneen Moolla, Senzo Mtshali, Veerle Msimang, Eftyxia Vardas, Jacqueline Weyer

**Affiliations:** Core Sequencing Facility, Centre for Emerging Zoonotic and Parasitic Diseases and Division of Public Health Surveillance and Response, National Institute for Communicable Diseases of the National Health Laboratory Service, Sandringham, South Africa; Lancet Laboratories, Johannesburg, South Africa; PathCare Laboratories, Cape Town, South Africa; AMPATH Laboratories, Centurion, South Africa; Department of Virology, University of the Witwatersrand, South Africa; Department of Molecular Pathology, University of Witwatersrand, South Africa; Department of Medical Virology, University of Pretoria, South Africa; Division of Medical Virology, Department of Pathology, Stellenbosch University; Division of Virology, University of the Free State, South Africa; Institute for Water and Wastewater Technology, Durban University of Technology, Durban, South Africa

**Keywords:** mpox, South Africa, multi-country outbreak

## Abstract

Mpox is an emerging viral infection and has since May 2022 been reported in a multi-country outbreak involving predominantly previously non-endemic countries. In addition, the number of cases of mpox is also rising in some endemic countries in Western and Central Africa. The disease is a notifiable medical condition in South Africa, with no cases reported before 2022. However, in 2022, coinciding with the peak of a multi-country mpox outbreak, five mpox cases were diagnosed in the country. Genomic sequencing and analysis revealed the presence of a monkeypox virus variant, which was circulating during the multi-country outbreak, namely Clade IIb sub-lineage B.1.7 (hMPXV). After the fifth case was reported in 2022, there were no cases detected for the following 20 months. In May 2024, mpox cases were again detected in South Africa leading to concerns that the virus may have been silently transmitting in the country since 2022. Genomic sequencing and analysis for 22 mpox cases reported during May to September 2024, were compared with sequences obtained from mpox cases reported in South Africa in 2022, and cases that have been reported globally. It was found that the sequences of the 2024 cases analysed in this study, clustered with Clade IIb hMPXV and sub-lineage B.1.20 and B.1.6 sequences. The results indicate that the detection of mpox cases in South Africa was an extension of the ongoing multi-country outbreak. Detection of different sub-lineages of hMPXV indicates reintroductions of the virus in the country since 2022, with local transmission of Clade IIb B1.20 hMPXV.

## Introduction

Mpox (previously monkeypox) is a viral disease associated with a painful rash. The disease is a zoonosis reported in 11 Western and Central African countries where historically limited human-to-human transmission has been reported.^1^ The infection is caused by the monkeypox virus (*orthopoxvirus monkeypox, Orthopoxviridae*) (MPXV) and a smallpox-like skin rash characterizes human disease.^2^ In otherwise healthy individuals, mpox is mild and self-limiting with little requirement for medical intervention.^3^ Mpox may however present as severe and possibly be a fatal disease in immunocompromised individuals, those living with co-morbidities, young children, and possibly also in pregnant females.^3^

Before 2022, mpox was recognized as a rare disease in humans. However, the number of reports from some endemic countries has been steadily increasing since 2000, and particularly from Nigeria and the Democratic Republic of Congo (DRC).^1,4^ Human-to-human transmission of the MPXV was previously uncommon, but since 2017, mpox has increasingly been associated with significant outbreaks characterized by sustained human-to-human transmission.^4-6^ Since 2017, following a quiescence of nearly 40 years, mpox cases have been reported from an expansive geographic range of Nigeria with human-to-human transmission chains.^7,8^ Following the re-emergence of mpox within Nigeria, mpox has been diagnosed in travelers to Nigeria returning to Singapore, Israel, the United Kingdom, and the United States of America (USA), with limited human-to-human transmission noted in a few of these cases.^8^ This increase in reports in travelers led up to a multi-country outbreak involving 116 countries, mostly outside of the known endemic zones of the disease, resulting in more than 99 000 cases since May 2022 and by June 2024.^6^ During the multi-country outbreak, cases have predominantly been reported in males, and men who have sex with men (MSM), with skin-to-skin contact during sex as the most frequently reported mechanism for viral transmission, and frequent lack of travel history to known endemic countries for mpox indicating sustained local human-to-human transmission.^6^ The first peak of the multi-country outbreak was reported in August 2022, followed by a significant decrease in cases, but with continued transmission noted in several countries.^6^ Following an initial decline, a resurgence of cases has been noted specifically in Europe during the second half of 2023, with a fivefold increase in the number of cases reported compared to the first six months of the year.^9^ The most notable resurgence was reported in Spain, Portugal, Germany, the United Kingdom, and France.^9^ Since 2022, an increased number of cases have also been reported from the DRC, with more than 1,000 mpox cases being laboratory-confirmed during the first six months of 2024, and evidence of human-to-human transmission.^1,4-6^ From July to August 2024, mpox associated with Clade Ib hMPXV has spread from the DRC to Burundi, Rwanda, Kenya and Uganda.^6^

Two phylogenetic branches of MPXV have been recognized based on geographic distribution and genomic characteristics, namely Clade I (previously Congo Basin Clade) and Clade II (previously West African Clade).^10,11^ Two distinct subclades are described for each of the clades.^11,12^ Clade Ia and Clade IIa have been primarily reported as zoonoses with limited human-to-human transmission, typically occurring in household clusters.^1,4,5^ Human mpox due to Clade IIa occurred in other parts of the world through the importation of African rodents into the USA in 2003 and by travelers from Africa but without evident human-to-human transmission.^1^ Since 2022, Clade Ib and Clade IIb viruses have been detected and have been associated with human-to-human transmission.^12^ Clade Ib viruses were initially noted in sex workers in the DRC in 2023.^12^ Since July 2024, cases of Clade Ib-associated mpox have been reported outside of the DRC.^6,13^ The emergence of Clade IIb was driven through human-to-human transmission of the virus in Nigeria with genetic analysis and modelling indicating that the virus may have been circulating undetected in the country since 2014.^14^ The divergent MPXV was subsequently denoted as hMPXV given the lack of zoonotic transmission events associated with its spread.^15^ Since May 2022, Clade IIb lineage B.1 has been disseminated through human-to-human transmission in a multi-country outbreak which in July 2024 is still ongoing.^6,16^ During the multi-country outbreak several sub-lineages of Clade IIb lineage B.1 have evolved, with the impact on phenotype not as yet understood.^14^

From June to August 2022, five cases of mpox were reported in South Africa (Source: NICD/NHLS). Histories obtained from cases were in keeping with possible exposure to persons from countries known to have mpox outbreaks, either directly through travel or through contact with international travelers (Source: NICD/NHLS). Two near-complete genomes could be generated from available clinical samples which were identified as Clade IIb lineage B.1.7.^17^ Whilst no cases were reported during 2023 in South Africa, cases have again been detected since 10 May 2024, with a total of 25 cases confirmed by 15 September 2024. Among these cases, only one reported an international travel history. Here we report on the findings of the genomic sequencing from clinical samples associated with confirmed mpox cases detected from May to September 2024 in South Africa.

## Methods

### Cases and ethics

The viral genomes derived from 22 cases of mpox diagnosed in South Africa during May and September 2024 and reported from the Gauteng, KwaZulu-Natal, and Western Cape provinces were included in the study. Details of the cases included in this investigation are summarized in Table 1. The Human Research Ethics Committee of the University of the Witwatersrand approved the analysis and reporting of these data - reference number: M210752 and protocol title: Essential communicable disease surveillance and outbreak investigation activities of the National Institute for Communicable Diseases.

**Table 1:** Data for mpox cases included in this study.

| **NICD laboratory**<br>**reference number** | **Date of diagnosis** | **City/town and Province where the case was identified** |
| --- | --- | --- |
| **SVPL 164/24** | 10 May 2024 | Johannesburg, Gauteng Province |
| **SVPL 175/24** | 23 May 2024 | Durban, KwaZulu-Natal Province |
| **SVPL 180/24** | 31 May 2024 | Durban, KwaZulu-Natal Province |
| **SVPL 189/24** | 1 June 2024 | Durban, KwaZulu-Natal Province |
| **SVPL 193/24** | 6 June 2024 | Johannesburg, Gauteng Province |
| **SVPL 201/24** | 10 June 2024 | Johannesburg, Gauteng Province |
| **SVPL 211/24** | 14 June 2024 | Durban, KwaZulu-Natal Province |
| **SVPL 221/24** | 14 June 2024 | Johannesburg, Gauteng Province |
| **SVPL 231/24** | 10 June 2024 | Cape Town, Western Cape Province |
| **SVPL 238/24** | 14 June 2024 | Tshwane, Gauteng Province |
| **SVPL 241/24** | 18 June 2024 | Durban, KwaZulu-Natal Province |
| **SVPL 273/24** | 21 June 2024 | Tshwane, Gauteng Province |
| **SVPL 279/24** | 21 June 2024 | Johannesburg, Gauteng Province |
| **SVPL 336/24** | 28 June 2024 | Johannesburg, Gauteng Province |
| **SVPL 346/24** | 28 June 2024 | Durban, KwaZulu-Natal Province |
| **SVPL 360/24** | 28 June 2024 | Tshwane, Gauteng Province |
| **SVPL 361/24** | 28 June 2024 | Fochville, Gauteng Province |
| **SVPL 413/24** | 25 June 2024 | Nquthu, KwaZulu-Natal Province |
| **SVPL 483/24** | 7 June 2024 | Nigel, Gauteng Province |
| **SVPL 737/24** | 26 July 2024 | Adams Mission, KwaZulu-Natal Province |
| **SVPL 767/24** | 7 August 2024 | Johannesburg, Gauteng Province |
| **SVPL 1015/24** | 10 September 2024 | Cape Town, Western Cape Province |

### DNA extraction and preparation

Dry swabs collected from skin lesions or scabs collected from lesions were vortexed or homogenized in 400µl phosphate-buffered saline. 140µl of the sample suspension was used to extract total viral nucleic acid using the QIACube QIAamp viral RNA mini kit (Qiagen, Valencia, CA, USA) using the QIACube extraction instrument (Qiagen, USA). Some samples were submitted to the NICD from private laboratories as nucleic acid extracts and were used without further manipulation. Random DNA amplification was performed by adding 5µl DNA to 40µl final reaction mix (5µl of 10x Expand High Fidelity buffer with 15 mM MgCl_2_, 1µl of 10mM dNTP, 1µl each of 100 µM primers Sol-PrimerA (5′-GTTTCCCACTGGAGGATA-N9-3′) and Sol-PrimerB (5′-GTTTCCCACTGGAGGATA-3′) ^18^ and 0.8 µl Expand High Fidelity enzyme mix (Roche, Basel, Switzerland). Reaction conditions for the PCR were: 94 °C for 2 min; 25 cycles of 94 °C for 30 s, 50 °C for 45 s, and 72 °C for 60 s, followed by 72 °C for 5 min. Targeted amplification for samples SVPL 737/24, SVPL 767/24, SVPL 201/24, SVPL 221/24, SVPL 241/24, SVPL 273/24, SVPL 279/24, SVPL 346/24, SVPL 413/24, SVPL 483/24, SVPL 211/24, SVPL 238/24, SVPL 1015/24 were carried out using iMap MPXV primers (2.5µl of 10x Expand High Fidelity buffer with 15 mM MgCl_2_, 0.5µl of 10mM dNTP, 3.6µl each of 10µM pool1 or pool2 primers, 0.4 µl Expand High Fidelity enzyme mix and 5 µl template in a total volume of 25 µl). Reaction conditions for the PCR were: 94 °C for 3 min; 35 cycles of 94 °C for 15 s and 63 °C for 5 min.^19^ The quantity and quality of the amplified DNA were analyzed using a Qubit fluorimeter (ThermoFisher, USA), as prescribed by the manufacturer. Amplified viral DNA was submitted to the NICD Sequencing Facility for whole-genome sequencing.

### Next-generation sequencing and analysis

The randomly amplified DNA was converted into Illumina sequence libraries using the Illumina DNA prep Kit (Illumina, San Diego, USA) according to the manufacturer’s instructions. Libraries were processed on an Illumina NextSeq 2000 (Illumina, San Diego, USA) to obtain ∼100 million reads (2×150bp). FASTQ files of the sequence data were trimmed for poor quality and adapter content using Trim Galore^20^ and aligned to the human genome reference sequence (GRCh38) using Bowtie2 version 2.4.2 in “sensitive” mode.^21^ SAMtools version 1.18 was used to convert the SAM alignment file to a BAM file, to label unmapped paired reads (-f 12 -F 256) and sort the BAM file.^22^ SAMToFastq version 3.0 was used to extract the unmapped reads from the BAM file.^23^ Unmapped reads were aligned to the MPXV-M5312 HM12 Rivers sequence (NC_063383.1) using Bowtie2 in “very-sensitive” model.^21^ SAMtools was used for SAM-to-BAM conversion, BAM sorting and indexing. SAMtools consensus in Bayesian consensus mode was used to extract a reference-guided assembly. Assemblies were submitted to NextClade version 3.8.2 and queried against the MPXV and hMPXV datasets for the initial clade assignment.^24^ The phylogeny produced by NextClade was used to collect sequences curated by the NextClade community, including genomic sequences that represent known MPXV and hMPXV lineages from the NCBI GenBank as of 19th April 2024. Additional Clade I MPXV genome sequences from the 2023-2024 DRC outbreak were collected from GISAID.^25,26^ The collection of sequences including samples from this study (see Table S1) with greater than 75% genome coverage was aligned using MAFFT version 7.487 in auto mode, and a phylogeny was produced using IQTREE v2.2.2.7 with 1000 bootstraps and automatic model selection.^27,28^

## Results

The coverage for the different samples varied, but near-complete genomes could be generated for phylogenetic analysis for eight of the twenty-two datasets (Table 2). With NextClade clade assignment, 21 sequences were clustered as hMPXV Clade IIb and the B.1.20 sub-lineage, and one as hMPXV Clade IIb and the B.1.6 sub-lineage (Figure 1).

**Table 2:** Next-generation sequence data mapping statistics for the data sets generated in this study.

| **Sample** | **Total number of reads** | **Number of reads mapping to reference\*** | **Depth of coverage against reference \*** | **Total genome size (bp)\*** | **Number of ambiguous bases** |
| --- | --- | --- | --- | --- | --- |
| SVPL 164/24** | 102,940,196 | 290,288 | 232 | 197,160 | 14 |
| SVPL 175/24** | 101,135,557 | 21,048 | 16 | 197,247 | 13,801 |
| SVPL 180/24** | 110,925,480 | 163,170 | 117 | 197,171 | 24 |
| SVPL 189/24** | 101,518,674 | 594,882 | 432 | 197,182 | 3 |
| SVPL 193/24** | 68,892,812 | 141,912 | 92 | 197,172 | 15 |
| SVPL 201/24** | 11,263,211 | 8,476,826 | 6,728 | 197,245 | 4,667 |
| SVPL 211/24** | 6,687,404 | 6,160,848 | 4,758 | 197,146 | 21,154 |
| SVPL 221/24 | 10,598,982 | 10,138,612 | 7,398 | 197,141 | 72,503 |
| SVPL 231/24** | 64,768,202 | 4,198,854 | 3,068 | 197,161 | 6 |
| SVPL 238/24** | 8,405,382 | 7,971,196 | 6,065 | 197,260 | 18,905 |
| SVPL 241/24** | 8,786,700 | 8,201,760 | 6,115 | 197,371 | 37,852 |
| SVPL 273/24 | 6,588,999 | 5,921,762 | 4,424 | 197,787 | 50,162 |
| SVPL 279/24 | 7,839,556 | 7,403,048 | 5,576 | 196,935 | 51,358 |
| SVPL 336/24 | 133,978,937 | 720 | 0,5 | 197,115 | 156,587 |
| SVPL 346/24 | 9,094,213 | 8,303,544 | 5,977 | 197,453 | 60,366 |
| SVPL 360/24** | 118,272,973 | 287,286 | 217 | 197,159 | 50 |
| SVPL 361/24** | 119,011,468 | 120,242 | 89 | 197,172 | 45 |
| SVPL 413/24 | 7,683,749 | 7,284,056 | 5,451 | 197,305 | 145,018 |
| SVPL 483/24 | 9,180,317 | 8,756,676 | 6,014 | 196,852 | 71,894 |
| SVPL 737/24** | 6,741,734 | 6,607,100 | 4,658 | 197,181 | 19,169 |
| SVPL 767/24 | 2,499,918 | 2,359,954 | 1,634 | 197,108 | 76,859 |
| SVPL 1015/24 | 5,722,639 | 5,066,138 | 3,746 | 197,296 | 65,033 |

**Figure 1.**
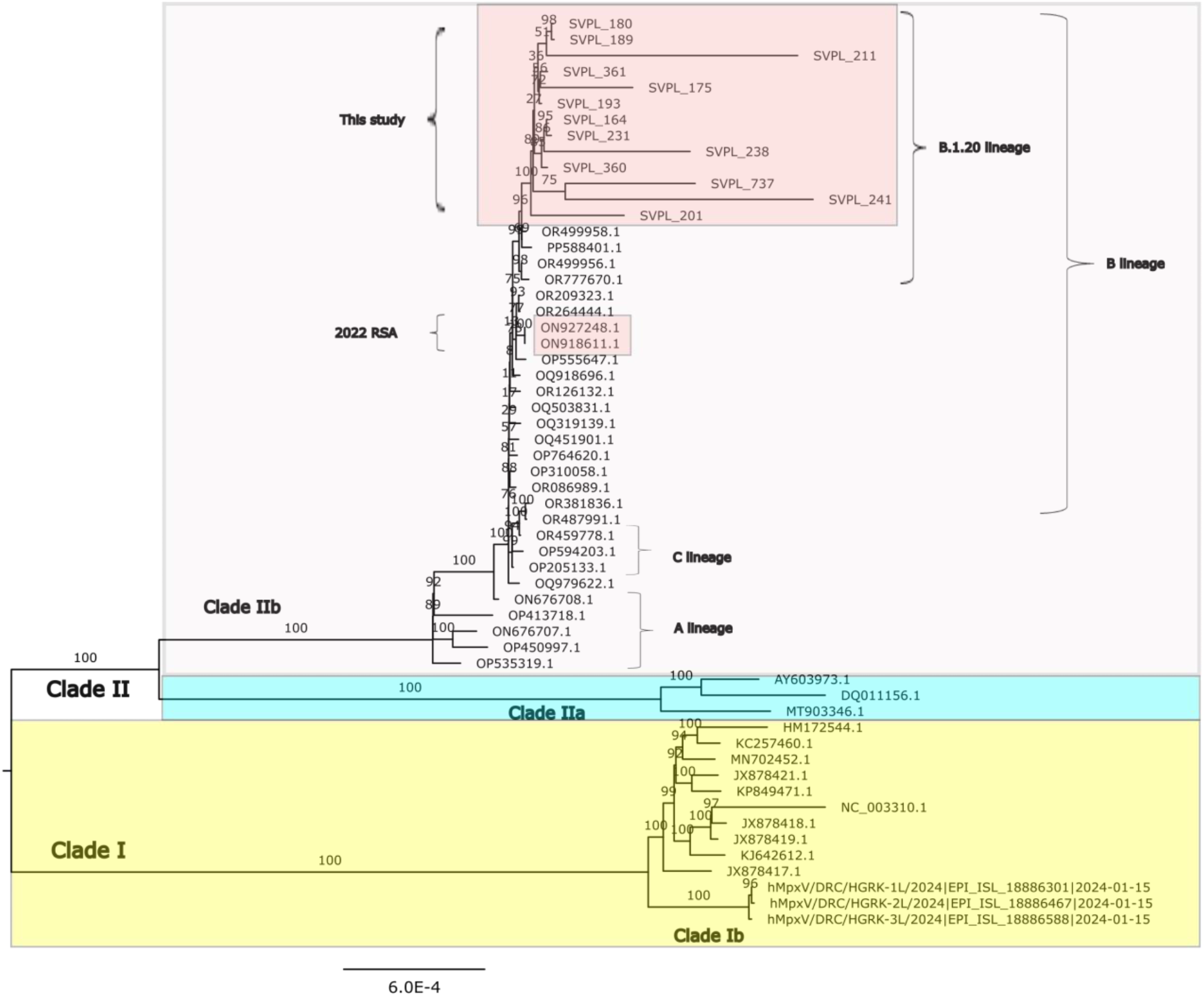
Maximum likelihood IQTREE v2.2.2.7 phylogeny of selected MPXV and hMPXV genomes produced with 1000 bootstraps and automatic model selection. Selected genomes from Nextstarin were included to encompass all known clades and most lineages to demonstrate the general topography of MPXV and hMPXV phylogeny. Genomes from this study with greater than 75% genome coverage were included. All hMPXV genomes for cases without travel history from the 2024, clustered with genomes from the B.1.20 sublineage based on Nexstrain lineage assignment (see Table S1).

## Discussion

During the multi-country mpox outbreak cases have been associated with Clade IIb, lineage B.1.^16^ This lineage has been widely detected across countries and continents, and initially did not allow for the tracking of cases geographically, but providing an indication of community transmission and continued international spread through travel.^16^ Subsequently, microevolution of the virus as a result of sustained human-to-human spread has however resulted in the detection of several sublineages of Clade IIb, lineage B.1. Mpox cases in South Africa in 2022, with available sequences, clustered with B.1 sequences detected internationally in 2022, linking the cases to the multi-country outbreak. With a growing number of genomic sequences available, these sequences were subsequently further classified as sublineage B.1.7.^29^ All the cases from 2024 without travel history, clustered with Clade IIb lineage B.1 and could further be assigned to sub-lineage B.1.20. One case with travel history to Peru prior to onset of illness, clustered with Clade IIb lineage B.1 and sub-lineage B.1.6.

The earliest genomic data available for sub-lineage B.1.20 relates to a case of mpox detected in the US in September 2022, with the largest number of B.1.20 sequences reported from the US since.^29^ The sub-lineage was subsequently detected in Germany in July 2023 and Portugal in December 2023.^29^ Likewise, sub-lineage B.1.6 was reported since June 2022, initially from Peru and subsequently also from Colombia, Spain, the United Kingdom and the US.^29^

From results reported here, the hypothesis is that mpox was introduced to South Africa through a traveler from the USA, Germany or Portugal, followed by sustained local transmission. The detection of cases associated with exposure during international travel, highlights the continued threat of introduction whilst mpox outbreaks are reported beyond South Africa’s borders.

## Conclusion

The occurrence of mpox in South Africa in 2022, and May – September 2024 have been an extension of the ongoing multi-country outbreak as demonstrated through phylogenetic analysis of near-complete hMPXV genomes. The detection of a different sublineage of Clade IIb B.1 in 2024 than what was detected in 2022, supports the hypothesis of a reintroduction of mpox in South Africa, possibly through a single introduction followed by local transmission. As mpox cases continue to be reported in the context of the multi-country outbreak, albeit at an overall lower level, the risk of international spread of the disease, causing outbreaks in new localities remains. In addition, the increase in detection of Clade Ib hMPXV in particular, from the DRC with spread to Eastern Africa in July 2024, poses another threat for the further spread of the virus. Apart from morbidity and mortality associated with the mpox outbreaks, the concern is that sustained and continued human-to-human transmission of hMPXVs will provide the evolutionary pressure which may result in possibly more transmissible or more virulent variants of the virus and presenting future public health challenges. Continued vigilance for mpox in aid of rapid public health responses which may interrupt chains of transmission is critical. In addition, monitoring of viral variants aids in the understanding of the epidemiology of the disease, and monitoring for genomic changes of the virus are required investments for preparedness for possible future threats posed by mpox.

## Author contributions

JF Methodology, formal analysis, data curation, writing - original draft

AAG Investigation, methodology, formal analysis, writing - original draft

AG, DG, KH, JM, KG, NG, AI, CLR, TM, DM, MM, NM, SM, VM, EV Investigation, methodology

JW Conceptualization, methodology, investigation, writing of original draft All authors contributed to the editing and reviewing of the draft.

## Acknowledgements

The authors would like to acknowledge the contributions of the staff of the Core Sequencing Facility at the NICD/NHLS in the processing of sequencing samples, which made this study possible.

## Data availability

Complete or near-complete genomes (i.e. sequences will less than 50 missing bases) were uploaded to the NCBI. The lab-identifier and corresponding GenBank ID of submitted genomes are: SVPL_164 (PQ141466), SVPL_180 (PQ141467), SVPL_189 (PQ141468), SVPL_193 (PQ141469), SVPL_231 (PQ141470), SVPL_360 (PQ213333), and SVPL_361 (PQ213334). These genomes and raw sequence reads were submitted under the BioProject number PRJNA1142891.

---

**Table S1.**
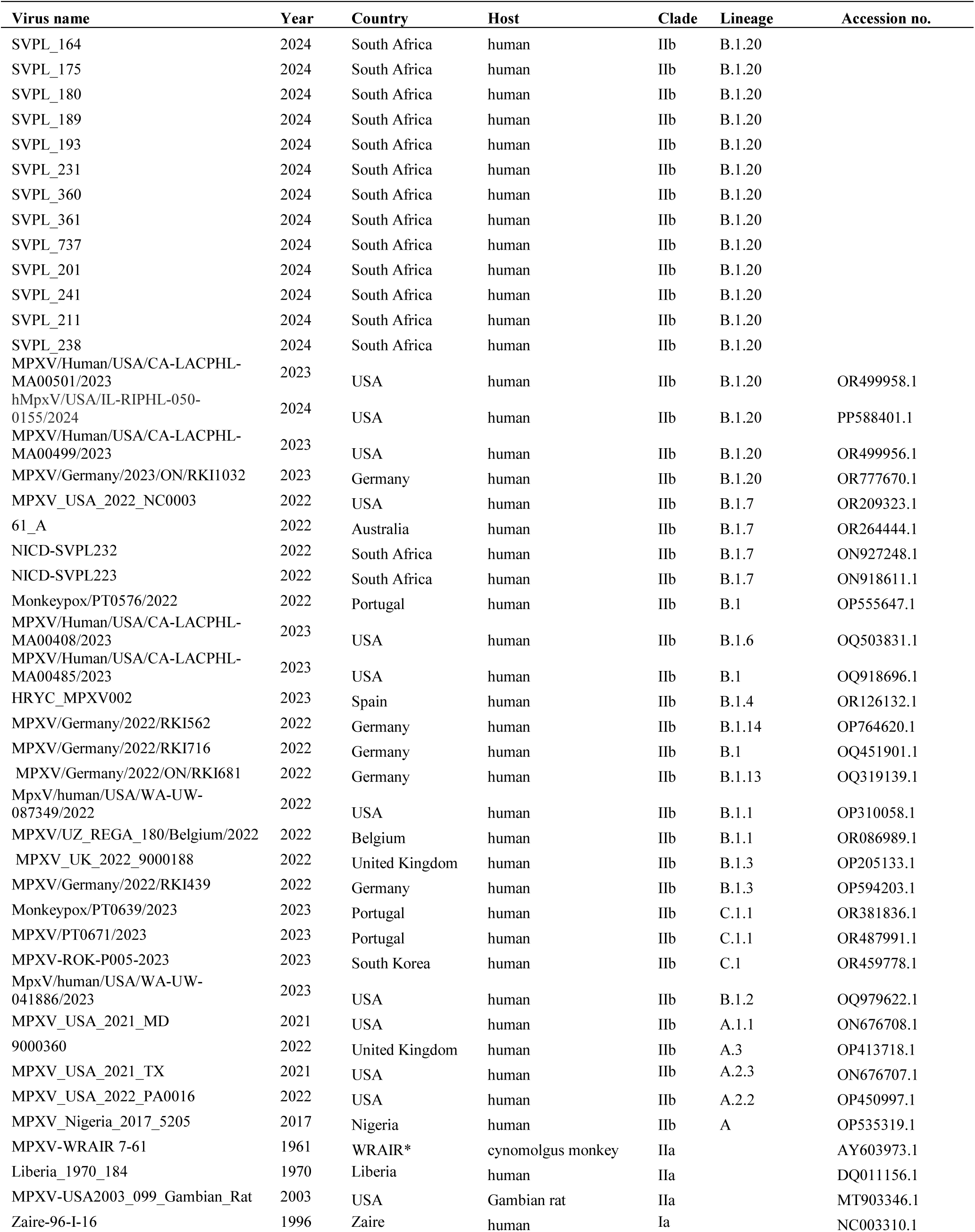
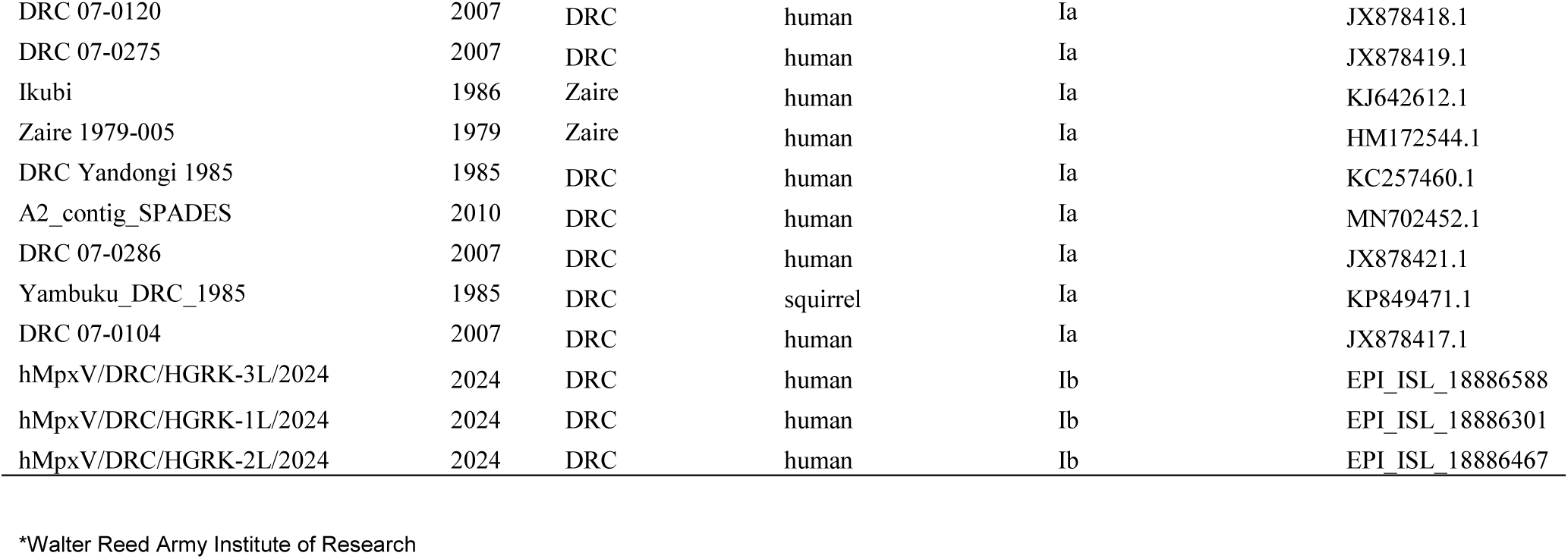
Details for sequences included in the phylogenetic tree.

